# Keeping common species common: the role of future climate refugia in species conservation

**DOI:** 10.1101/2024.07.19.602245

**Authors:** Chiara Serafini, Nina Luisa Santostasi, Daniele Canestrelli, Gentile Francesco Ficetola, Luigi Maiorano

## Abstract

Climate change is one of the most important challenges for biodiversity conservation. Species may respond to changing climates by moving, adapting, and/or adjusting. The move response is the easiest and quickest as it does not imply any evolutionary and/or physiological response.

However, moving in space to track changing climate is not an option for species with restricted movement capacities (e.g., many amphibians) or species endemic to islands. Therefore, the impact of climate change on these species is potentially dramatic, even when they are currently widespread and least concern. Planning for the conservation of these species in a global change context requires a proactive approach, with the identification of climatic refugia, i.e., areas climatically suitable for a given species under both current climate and future scenarios.

Here, we demonstrated our approach considering the *Hyla sarda*, an amphibian endemic of the islands of Sardinia and Corsica, currently widespread in its range, and Least Concern according to the IUCN Red List. We calibrated an SDM for the species focusing on Sardinia and projected it into the future, identifying all areas that can act as future climatic refugia. We also evaluated the coverage of the refugia by the existing protected areas.

According to our results, *Hyla sarda* will experience a significant restriction of its distribution range due to projected climate changes, with small and highly fragmented climatic refugia mostly located outside of existing protected areas. Our findings highlight the importance of considering common species in global change studies. All our conservation strategies should be more proactive if we want to conserve common species before they become rare.

## 1. INTRODUCTION

Climate change is projected to quickly become the major driver of biodiversity loss (Bellard et al., 2012), with species that are already responding by adjusting (Ficetola & Maiorano 2016), adapting (McCulloch & Waters 2023), moving (Rubenstein et al. 2023), or going extinct (Holzmann et al. 2023). The move response is certainly the most widely studied and, biologically and ecologically, the simplest as it does not imply any physiological and/or evolutionary response (Parmesan 2006). Basically, species shift their distribution to track the changing climate and can see a recession (with a range restriction), an expansion, a shift, or a combination of the three (Lenoir & Svenning 2015).

Clearly, to expand their range or to shift their distribution species must rely on movement. However, many species have very limited dispersal capabilities (e.g., many amphibians and many reptiles) or simply have no real possibility of moving (e.g., many species endemic to islands). When these two conditions are combined, for example for the rich herptile fauna of the Mediterranean islands, even species that currently are considered least concern and widespread can become endangered by climate change.

In this context, a proactive conservation approach is absolutely needed. Protected areas are the core of our in situ conservation strategies, and are expected to mitigate the effects of global change on biodiversity (Hannah et al., 2002). A huge share of species in both amphibians and reptiles is currently protected at the global level (Mi et al. 2023), but if we focus on the European continent, protected areas do not cover amphibians better than what would be expected by chance (Sánchez-Fernández & Abellán 2015), and the problem is even more important for range-restricted species which often are endemic to Mediterranean islands (Nori et al.,2015).

Furthermore, climate change is already acting as the main threat for 39% of the amphibians, followed by habitat loss (for 37% of the amphibians), and the combined effect of both is projected to increase over time (Luedtke et al. 2023). Being ectothermic, amphibians may be particularly vulnerable to climate change (Pimm et al., 2014), and even widespread species currently considered least concern may risk a collapse due to the changing climate.

It is therefore extremely important to identify future climate refugia, areas whose climate suitability is projected to remain stable in time and that should act as the cornerstone of any onsite conservation strategy. These climate refugia should be included in the network of protected areas to buffer land use changes (but see Maiorano et al. 2008), especially in the human-dominated Mediterranean basin.

Here we apply the idea of future climate refugia to the Sardinian tree frog (*Hyla sarda*), an amphibian endemic to Sardinia, Corsica, and the Tuscan archipelago (Italy and France). We focused our analyses on Sardinia, a Mediterranean island and a hotspot of endemism for endemic amphibians (Maiorano et al. 2013). The Sardinian tree frog is a widespread species, occurring on the entire island, and it can be present at very different elevations, although it is more abundant along the coasts (Lanza et al., 2007). It is classified as “Least Concern” in the IUCN Red List, given its relatively wide distribution, the high tolerance to habitat modification, and the presumed large total population (https://www.iucnredlist.org).

We calibrated a multi-scale correlative species distribution model (SDM) considering the current distribution of the tree frog in Sardinia and we projected the SDM in time under different future climate scenarios to explore the following research questions: a) is climate change going to be a factor in the conservation of a common, widespread species which cannot move outside of the current distribution range? b) can we identify future climate refugia for these species? c) are existing protected areas effectively covering the future climatic refugia?

## 2. METHODS

### 2.1 Study area

We focused all our analyses on Sardinia, the second biggest island in the Mediterranean basin (24,100 km^2^). Sardinia is mainly mountainous and hilly in its eastern part (max elevation 1835 m), with two plains located in the western portion of the island. The climate is typically Mediterranean (dry and hot summers, relatively rainy and mild winters), with small inlets of temperate climate corresponding to the highest elevations.

### 2.2 Species occurrences and background data

We downloaded occurrence data for the Sardinian tree frog from two sources: GBIF (Global Biodiversity Information Facility, https://www.gbif.org/) and iNaturalist (https://www.inaturalist.org/). We removed duplicates (same coordinates, same dates) and unreliable records (Zizka et al., 2019) and retained in the analysis only the occurrences sampled after 2000 and with a positional accuracy ≤ 1km. Overall, the final dataset that we used for model calibration comprised 181 occurrences distributed all over Sardinia (Figure 1).

**Figure 1.**
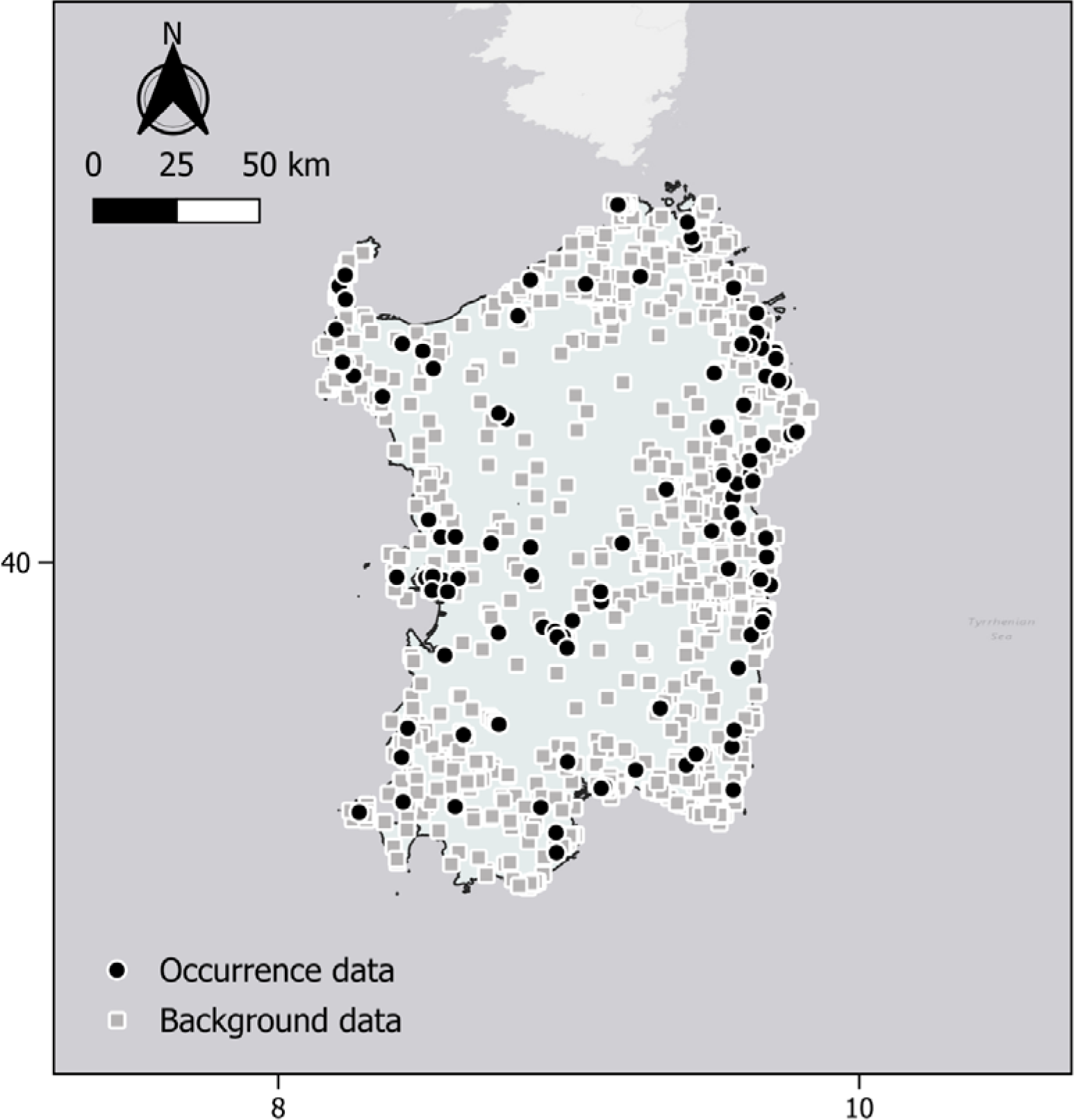
The occurrence data and the background data used to calibrate the model for the Sardinian tree frog (*Hyla sarda*) in Sardinia (Italy).

To minimize the biases commonly present in online repositories of occurrence data (Hortal et al., 2015; Garcia-Rosello et al., 2023), we calibrated our SDM considering a target group approach (Ranc et al., 2017; Barber et al., 2022). Hence, we selected as background points all the occurrences of amphibians and reptiles for Sardinia from GBIF and iNaturalist, assuming that they shared approximately the same sampling bias of the occurrence points. We cleaned background points following the same procedure described for the occurrence data. Overall, the background dataset comprised 1824 points (Figure 1).

### 2.3 Predictor variables

To calibrate the multiscale SDM, we considered a set of large-scale bioclimatic variables plus a local-scale water availability variable. We obtained bioclimatic variables for the current time frame (average from 1981 to 2010) and future projections (average from 2041 to 2070, hereafter 2050; average from 2071 to 2100, hereafter 2100) from CHELSA version 2.1 (Karger et al., 2017). For the future projections, following the Intersectoral Impact Model Intercomparison Project recommendations, we selected five different General Circulation Models (GCMs: GFDL-ESM4, IPSL-CM6A-LR, MPI-ESM1-2-HR, MRI-ESM2-0, UKESM1-0-LL) and three different Shared Socio- economic Pathways from the CMIP6 scenarios (ssp 1-2.6, ssp3-7.0, and ssp 5-8.5). From the initial set of 19 bioclimatic variables, we excluded isothermality and the annual range of air temperature since they are built as linear combinations of other bioclimatic variables.

Since the Sardinian tree frog is strictly dependent on water to complete its biological cycle and it seldom leaves the water edges, even outside the reproductive season (Lanza et al., 2007), we calculated a local scale water availability index. Water availability at each point is strictly dependent on topography and precipitation. Therefore, to map potential ponding areas, we first built a Topographic Wetness Index considering the effect of local topography on runoff and accumulation.

TWI is a function of both the upstream contributing area and the slope and was obtained from a 10m resolution digital elevation model (Tarquini et al., 2023) applying the following formula:

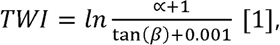

where ex represented flow accumulation, and f3 represented the local slope in radians. This formula is a correction of the original formula (Beven and Kirkby 1979) that generates finite values when tan(f3) = 0. All negative values were set to 0. All calculations were performed in ArcGIS pro (ESRI ©).

The TWI was used to calculate the Precipitation Topographic Wetness Index (PTWI), an effective indicator of the presence of wetlands (Hu et al., 2017). PTWI was calculated following Hu et al., 2017:

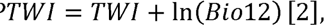

where Bio12 is the mean annual precipitation, downloaded from CHELSA version 2.1 (Karger et al., 2017). TWI was assumed to remain unchanged over time while Bio12 was changed following different GCMs and scenarios (see above).

The Sardinian tree frog does not occur in large permanent water bodies and rivers, and we excluded all these areas from the study area using as a mask the layer of permanent waters obtained from the ESA Worldcover 2020 (Zanaga et al., 2021) at 10 m resolution.

To limit multicollinearity, we performed a Variance Inflation Factor (VIF) analysis with a threshold of VIF=3, obtaining a final set of 5 uncorrelated predictors (Table 1).

**Table 1:**
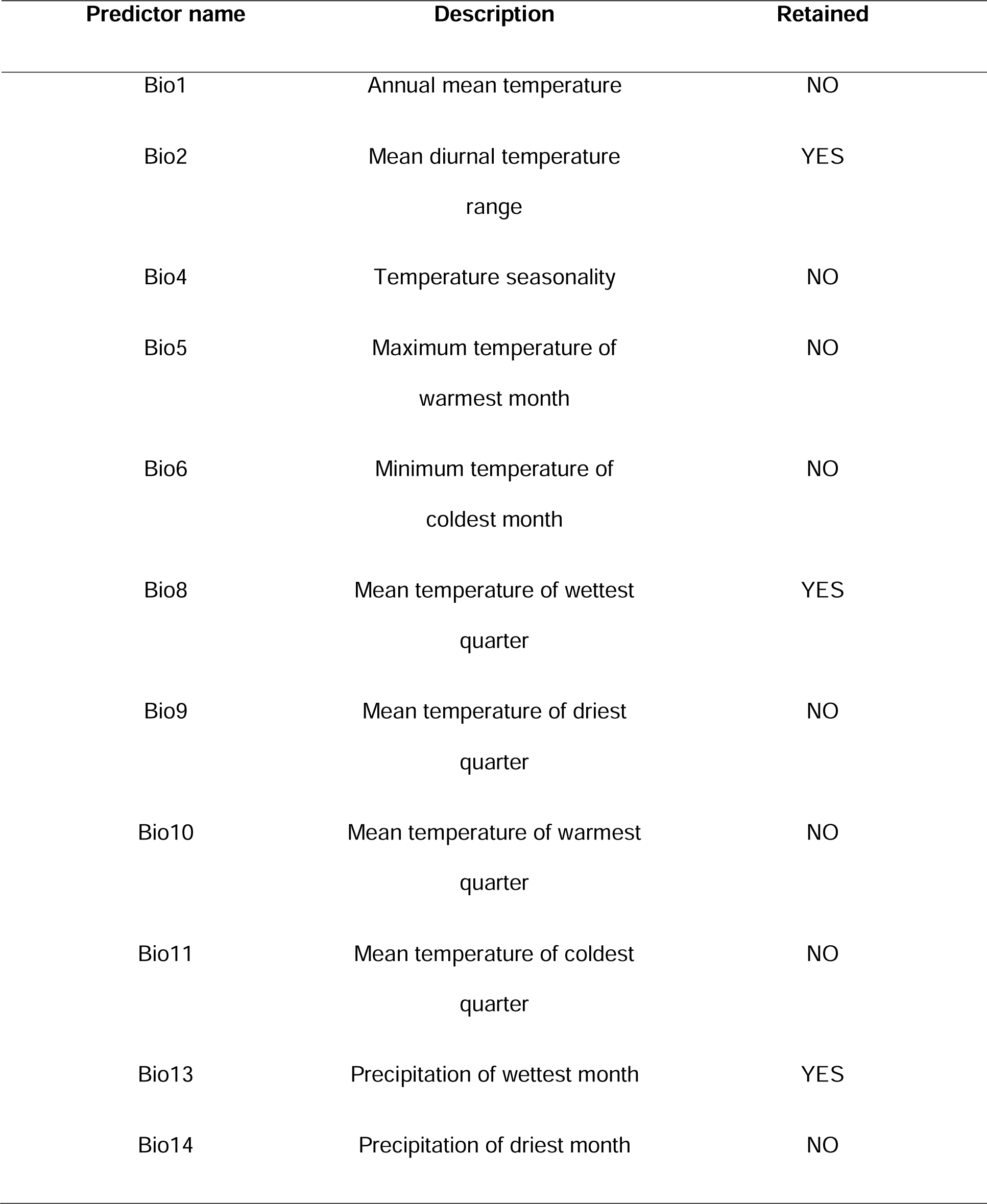
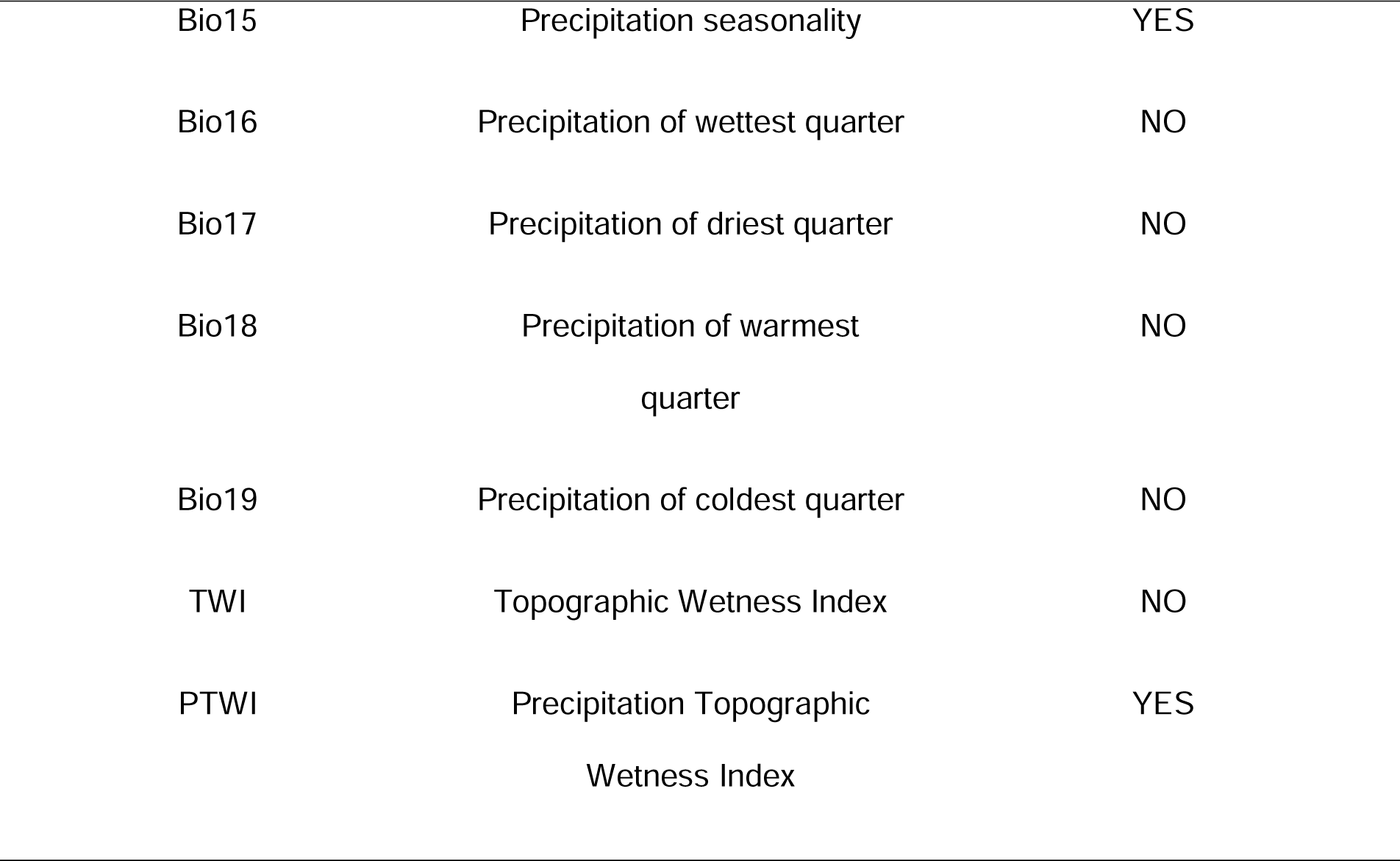
Predictor variables considered for modelling the distribution of *Hyla sarda* in Sardinia (Italy). The column “Retained” indicates whether the variable was used in model calibration or excluded to avoid of multicollinearity.

| Predictor name | Description | Retained |
| --- | --- | --- |
| Bio1 | Annual mean temperature | NO |
| Bio2 | Mean diurnal temperature<br>range | YES |
| Bio4 | Temperature seasonality | NO |
| Bio5 | Maximum temperature of<br>warmest month | NO |
| Bio6 | Minimum temperature of<br>coldest month | NO |
| Bio8 | Mean temperature of wettest<br>quarter | YES |
| Bio9 | Mean temperature of driest<br>quarter | NO |
| Bio10 | Mean temperature of warmest<br>quarter | NO |
| Bio11 | Mean temperature of coldest<br>quarter | NO |
| Bio13 | Precipitation of wettest month | YES |
| Bio14 | Precipitation of driest month | NO |
| Bio15 | Precipitation seasonality | YES |
| Bio16 | Precipitation of wettest quarter | NO |
| Bio17 | Precipitation of driest quarter | NO |
| Bio18 | Precipitation of warmest<br>quarter | NO |
| Bio19 | Precipitation of coldest quarter | NO |
| TWI | Topographic Wetness Index | NO |
| PTWI | Precipitation Topographic<br>Wetness Index | YES |

### 2.4 Species Distribution Model

We modelled the distribution of the Sardinian tree frog with Maxent, a machine-learning approach that estimates the most uniform distribution (maximum entropy) of the occurrences compared to the background locations given the constraints defined by the environmental data (Phillips et al., 2006). We chose Maxent since it has been shown to outperform other statistical techniques in different studies (e.g. Hernandez et al., 2006; Wisz et al., 2008), particularly when fine-tuned for the species/area considered (Valavi et al., 2021). We selected the best settings for our model considering a set of thirty alternative models with different combinations of feature classes and regularization multipliers and measuring the sample size corrected Akaike Information Criteria (AICc, Anderson & Burnam 2004). We performed all analyses using the “ENMeval” package in R version 4.3.2 (Muscarella et al., 2014; Kass et al., 2021). To evaluate model performance in prediction we used a 10-fold cross-validation and we calculated the area under the receiver operating characteristic curve (AUC). We tested the statistical significance of the AUC values against a null distribution of 100 expected AUC values based on random data, following the framework proposed by Bohl et al. (2019) and implemented in the “ENMeval” package.

We assessed variable importance considering percent contribution (Phillips et al., 2006) and we projected the model on current and future climate conditions. For each future socio-economic scenario, we calculated a final continuous model as the average among the results obtained with the five different GCMs.

### 2.5 Climatic refugia and gap analysis

To map future climatic refugia for the Sardinian tree frog we binarized all the continuous relative probability maps using the median probability of the training presences under the current climate as a threshold. Using the binary projections, we defined climatic refugia using two alternative approaches: a majority approach, in which an area is a climatic refugia if the majority of the GCMs (i.e., at least 3 GCMs out of 5) predict a presence for the species, and a consensus approach, in which an area is a climatic refugia only when all GCMs predict a presence for the species. We performed the same analyses for the three socio-economic scenarios, obtaining therefore alternative maps of climate refugia.

To evaluate whether the protected areas cover the future climatic refugia, we performed a gap analysis (sensu Maiorano et al. 2015) considering all existing protected areas of Sardinia, (obtained from www.protectedplanet.net).

## 3. RESULTS

### 3.1 Evaluation of Predictive Performance and variable importance

The best tuning settings according to the AICc values were the Hinge feature with a regularization multiplier equal to 2. The mean test AUC for the best model was 0.68, significantly better (p < 0.05) if compared to the mean test AUC obtained with random occurrences for the same study area and variables (0.53±0.01).

The Sardinian tree frog showed a positive response to increasing precipitation seasonality (BIO15), increasing PTWI, mean diurnal temperature range (BIO2) over 2 °C, and precipitation of the wettest month (BIO13) above 62 mm (Table 2; Fig. 2). The mean temperature of the wettest quarter (BIO8) was not important in predicting species presence.

**Figure 2.**
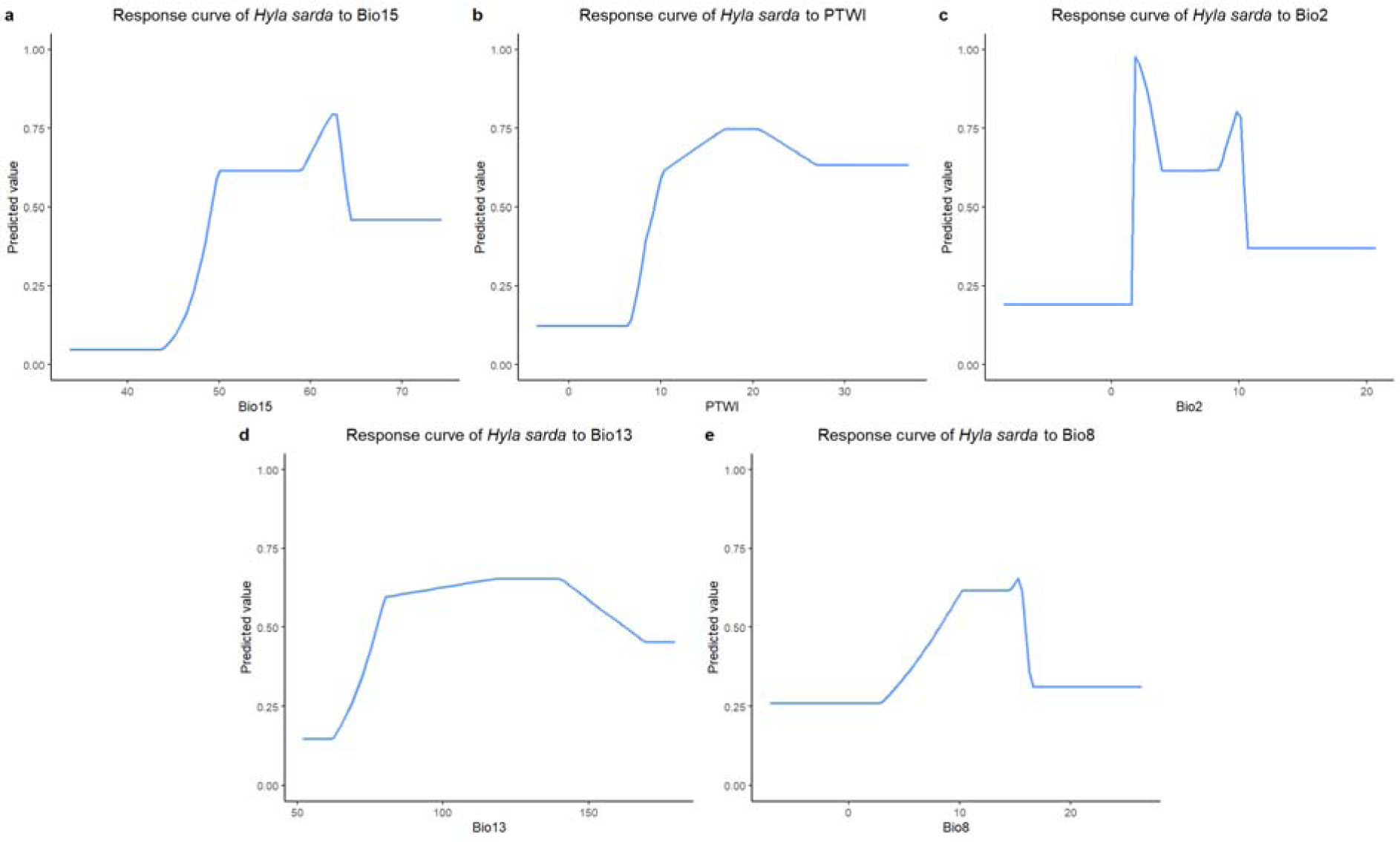
The response curves for the Sardinian tree frog model **a)** Precipitation seasonality (Bio15) **b)** Precipitation Topographic Wetness Index (PTWI) **c)** Mean diurnal temperature range (Bio2) **d)** Precipitation of the wettest month (Bio13) d) Mean temperature of the wettest quarter (Bio8).

**Table 2:** Variable importance measured as percent contribution.

| Predictor | Percent contribution |
| --- | --- |
| BIO15 | 35 |
| PTWI | 28 |
| BIO2 | 17 |
| BIO13 | 16 |
| BIO8 | 5 |

### 3.2 Current and future projections

Under current climate conditions, the probability of presence of the Sardinian tree frog is high over large areas of the island (Fig. 3), particularly in the western part, which is characterized by the highest values of precipitation seasonality and PTWI. Only the Limbara and Gennargentu mountains, corresponding to the highest elevations in Sardinia and located in the eastern portion of the study area, are completely unsuitable for species presence.

**Figure 3.**
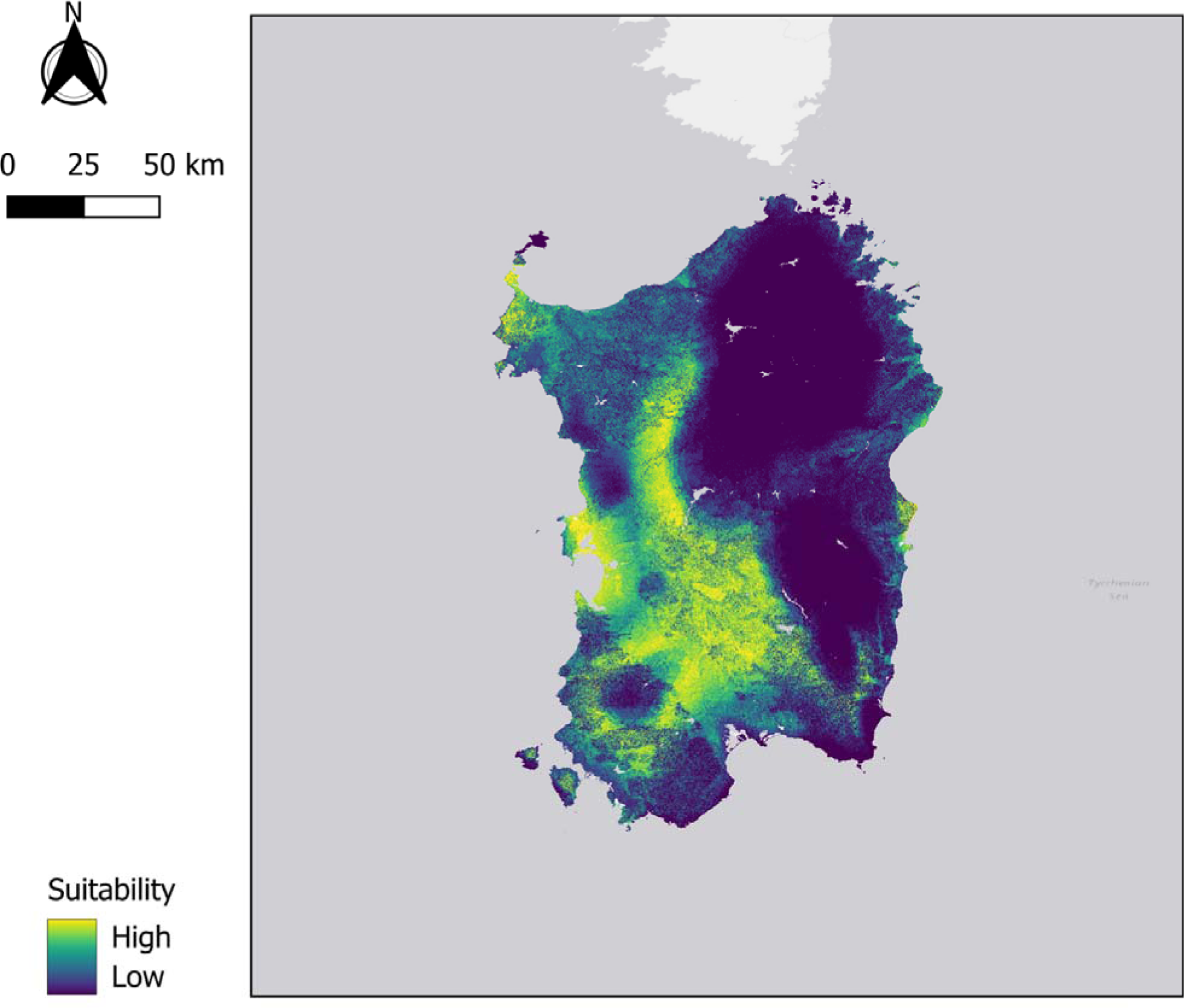
Current projection of the potential distribution of *Hyla sarda* in Sardinia (Italy).

Considering future climate scenarios (Fig. 4), the probability of presence for the species is always projected to probabilities much lower compared to the present, with most of the suitability being lost also in the western part of the island. In the best possible scenario, the ssp 1-2.6 for the 2050 time frame, the species is projected to lose up to 74% of its current potential distribution. The loss of suitable areas is projected to a maximum of 95% considering the ssp 5-8.5 scenario for the 2100 time frame.

**Figure 4.**
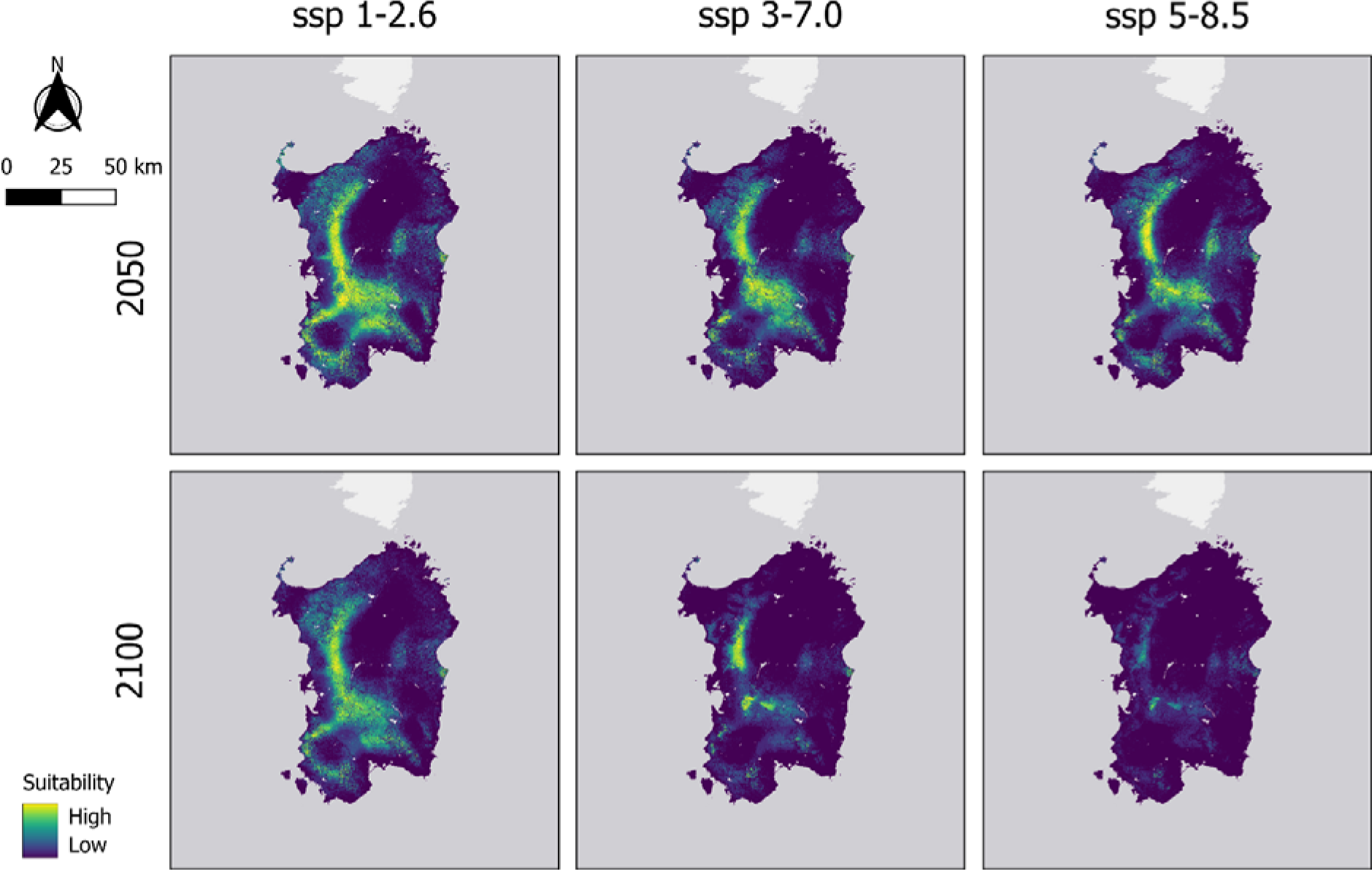
Potential distribution of the Sardinian tree frog for the 2050 and the 2100 timeframes under three different socio-economic scenarios: ssp 1-2.6, ssp 3-7.0, and ssp 5-8.5.

### 3.3 Climatic refugia and gap analysis

Going from mild climate change scenarios towards extreme scenarios, the areas that will act as climatic refugia are projected to decrease from 43 km^2^ (ssp 1-2.6) to 19 km^2^ (ssp 3-7.0) and, finally, to only 2 km^2^ (ssp 5-8.5) when considering the majority of GCMs. However, when we focus on consensus climate refugia (all the 5 GCMs in agreement), the refugia went from 8 km^2^ (ssp 1- 2.6) to 2 km^2^ (ssp 3-7.0), and finally to 0.02 km^2^ (ssp 5-8.5) (Figure 5). Whatever approach is considered, the refugia under milder climate scenarios are predicted to occur in the western part of Sardinia, but under extreme scenarios, the few and small refugia are almost exclusively located in the eastern part of the island (the Orosei Gulf).

**Figure 5.**
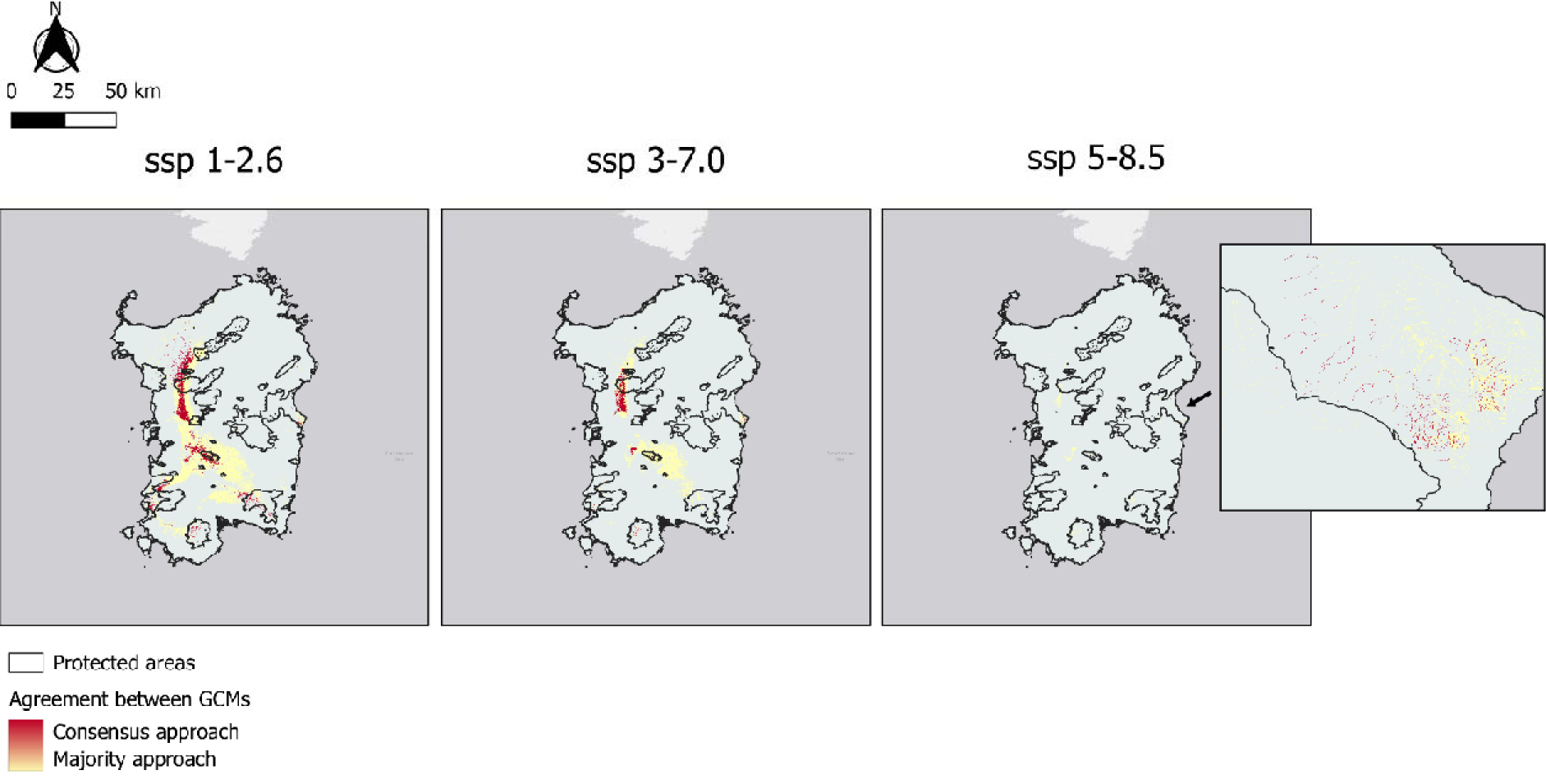
The climatic refugia under 3 different socio-economic scenarios (ssp 1-2.6, ssp3-7.0, ssp 5-8.5) following 2 approaches to determine the refugia: Majority approach (3 GCMs out of 5 in agreement), and Consensus approach (all the 5 GCMs in agreement).

Existing protected areas covered from 16% (majority approach) to 21% (consensus approach) of the refugia for the ssp 1-2.6, from 18% (majority approach) to 28% (consensus approach) of the refugia for the ssp 3-7.0, and from 31% (majority approach) to 88% (consensus approach) of the refugia for the ssp 5-8.5.

## 4. DISCUSSION

Climate change is expected to markedly impact biodiversity, potentially forcing species to adapt, adjust, move, or go extinct. Adapting or adjusting to the changing climate often implies a number of complex physiological and genetic responses (Parmesan et al., 2006), while the move response is often regarded as the easiest and quickest (Parmesan et al., 2006). Therefore, when the possibility of a shift in distribution is limited due to the movement capacities of a species (such as for amphibians) and/or to lack of available options (e.g., species living on islands), even widespread species currently not at risk of extinction can face important threats under future climate scenarios. Here, we propose a framework that can be used to identify potential future climatic refugia. We focused our case study on the Sardinian tree frog, an amphibian endemic of Sardinia, Corsica, and a few small islands in Tuscany. We found that, while the species is currently widely distributed, its future distribution is restricted especially under extreme climate change scenarios (e.g., the ssp 5- 8.5). The climatic refugia, i.e., those areas that will retain climatic suitability in time, are going to be small and fragmented, and current protected areas only cover a limited portion of these refugia.

It’s worth noting that the Sardinian tree frog is projected to lose a staggering 90% of its current distribution, making it a clear candidate to jump from Least Concern to Critically Endangered in the next 50 years under the ssp 5-8.5 scenario. While the ssp 5-8.5 is considered a worst-case scenario in the latest Coupled Model Intercomparison Project Phase 6 (CMIP6), it is the scenario that most closely matches the cumulative emissions today (Schwalm et al. 2020), and economic growth rates higher than today but still very plausible could make the ssp 5-8.5 35% more likely (Christensen et al. 2018). In this context, if we want to avoid losing another species in a conservation triage (Bottrill et al. 2008), it’s important that we plan today for the conservation of the next 50 to 100 years.

The Sardinian tree frog possibly represents only an example among many possibilities. Previous studies have clearly demonstrated that most amphibians living on islands in the western Mediterranean have a very limited capacity to shift their distribution under climate change (e.g., Enriquez-Urzelai et al., 2019). For all these species, the identification of future climatic refugia represents a clear priority for conservation, as they possibly represent the only option to preserve these species under climate change. While conservation actions are certainly urgent for rare and threatened species (Boyd et al., 2008), it is important to develop approaches to conservation planning focusing also on common species and their future.

